# Multilayer network analysis of miRNA and protein expression profiles in breast cancer patients

**DOI:** 10.1101/384768

**Authors:** Yang Zhang, Jiannan Chen, Dehua Wang, Weihui Cong, Bo Shiun Lai, Yi Zhao

**Affiliations:** Harbin Institute of Technology (Shenzhen), Shenzhen, Guangdong, 518055, China; Johns Hopkins University School of Medicine, Baltimore, Maryland 21205, United States

## Abstract

MiRNAs and proteins play important roles in different stages of tumor development and serve as biomarkers for the early diagnosis of cancer. A new algorithm that combines machine learning algorithms and multilayer complex network analysis is hereby proposed to explore the potential diagnostic values of miRNAs and proteins. XGBoost and random forest algorithms were employed to exclude unrelated miRNAs and proteins, and the most significant candidates were retained for the further analysis. Given these candidates’ possible functional relationships to one other, a multilayer complex network was constructed to identify miRNAs and proteins that could serve as biomarkers for breast cancer. Proteins and miRNAs that are nodes in the network were subsequently categorized into two network layers considering their distinct functions. Maximal information coefficient (MIC) was applied to assess intralayer and interlayer connection. The betweenness centrality was used as the first measurement of the importance of the nodes within each single layer. To further characterize the interlayer interaction between miRNAs and proteins, the degree of the nodes was chosen as the second measurement to map their signalling pathways. By combining these two measurements into one score and comparing the difference of the same candidate between normal tissue and cancer tissue, this novel multilayer network analysis could be applied to successfully identify molecules associated with breast cancer.

## Introduction

Breast cancer is the second leading cause of cancer death among women and results in millions of new cases every year [1]. Often assuming regulatory roles in eukaryotic cells, miRNAs are small, non-coding RNAs of roughly 20~22 nucleotides that can bind to and inhibit protein coding mRNAs [2]. The expression profiles of miRNAs are correlated with cancer type, stage, and other clinical variables [3]. Therefore, miRNA expression profiling could be a useful tool for cancer diagnosis and prognosis. MiRNAs play important roles in almost all aspects of cancer biology, including proliferation, apoptosis, tissue invasion, metastasis, and angiogenesis [4]. miRNAs also play important roles in toxicogenomics and may explain the relationship between toxicant exposure and tumorigenesis. Previous work has identified 63 miRNA genes shown to be epigenetically regulated in association with 21 diseases, including 11 cancer types [5].

Many proteins have known oncogenic properties that contribute to tumorigenesis. Therefore, proteomics data can also be used to study the characteristics and observe the presence of cancer [6], and used as biomarkers for breast cancer to provide targets for the early diagnosis and intervention [7].

Machine learning plays increasingly important roles in cancer diagnosis [8]. Algorithms such as Bayes, decision tree, and support vector machine, are widely used in the classification of breast cancer [9]. Deep learning methods like convolutional neural network are also prevalent in the analysis of biopsy images of cancer [10]. Previous studies have compared the performance of various statistical methods in classifying cancer based on Mass Spectrometry (MS) spectra. These methods encompass linear discriminant analysis, quadratic discriminant analysis, k-nearest neighbour classifier, bagging and boosting classification trees, support vector machine, and random forest (RF). It has been demonstrated that RF outperforms other methods in the analysis of MS data [11]. As a result, the RF algorithm was adopted for filtering miRNAs and proteins, thereby retaining the most relevant biomarkers. Furthermore, to reduce the chance of missing important biomarkers, two established ensemble learning methods -- random forests and XGBoost, were employed to complete the feature selection result.

Binding of miRNAs to mRNAs leads to destabilization or translational repression of the target mRNA, which in turn regulates the expression of protein. Multiple miRNAs and proteins known to be involved in different signalling pathways are deregulated in breast cancer. To improve understanding of interaction between miRNAs and cancer protein, multilayer networks consisting of protein and miRNA expression was constructed in this study. In the multilayer network, miRNAs and proteins are regarded as nodes in each layer. Both the MIC values between nodes within each layer and between two separate layers were computed to determine whether there exists intralayer and interlayer edges between any two nodes under a specific threshold of MIC. This model consists of multiple subsystems and multiple connectivity layers, allowing different dynamic processes to be coupled and improving our visual understanding of multilayer systems. In particular, this biological multilayer network model exhibits the interrelationship between the miRNA and protein, thereby studying their combined action on cancer at different scales and levels.

To better understand their roles in the context of biological networks, miRNA and protein expression profile networks were constructed for both normal and breast tissues. Furthermore, due to the large number of miRNAs and proteins, in order to prevent analysis process from interference of unrelated variables, random forest model and XGBoost were applied to filter miRNAs and proteins before establishing a multilayer network. The filtered molecules were used as nodes in the network. Both threshold and MIC values between every two nodes determined the final structure of the multilayer network. Comparing the betweenness centrality of the node between health control and patient samples could lead to the novel finding of miRNAs and proteins related to cancer.

## Materials and methods

### Process overview

Schematic representation of data processing and analysis is shown (Fig 1). XGBoost and random forest algorithms were employed for feature selection, and the results were used for the subsequent processing step. Subsequently, MIC value for any two nodes was calculated, so that the weight network of expression data can be obtained. By setting specific threshold, the MIC values are then converted into Boolean variables, resulting in a complex network without weights. Finally, score related to breast cancer of each node was computed and nodes were ranked by scores.

**Fig 1.**
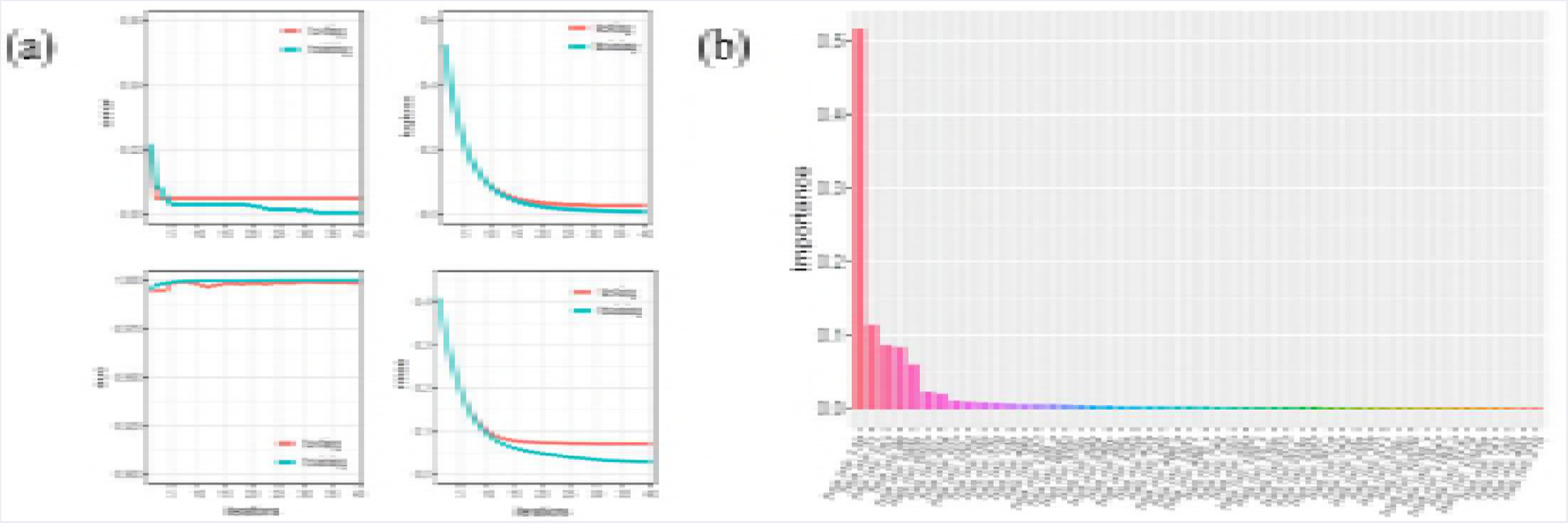
Schematic representation of data processing and analysis. Each icon denotes an analytical process. Icon 1 denotes data containing miRNAs and proteins. Icon 2 shows the feature selection process, which includes XGBoost and random forest algorithms. Icon 3 indicates calculation of the MIC value for every two nodes in the network, which represents the interaction between nodes. Icon 4 is the process that generates edges in the network by setting a specific threshold of MIC. Icon 5 represents the construction of a multilayer network. Icon 6 is the final step of analysis process that gives each node an importance score related to breast cancer.

### Data

Experimental data were collected from the Cancer Genome Atlas/TCGA (https://tcga-data.nci.nih.gov/tcga/dataAccessMatrix.htm). miRNA expression data consists of 1200 patient samples exploring expression level of 320 different miRNAs. Among them, 1096 cases are tumor tissues and the rest are normal tissues. Protein expression data consists of 925 patient samples investigating expression level of 199 candidate proteins. 882 cases are tumor tissues and the rest are normal tissues.

### Random forest algorithm and feature selection

Since random forest performed well on Mass Spectrometry spectra data, the same method was used for miRNA and protein expression profiles data [11]. Random forest [12], as one kind of ensemble learning method, in which each learning algorithm is a decision tree. Unlike in an ordinary decision tree, k attributes are first selected as candidate attributes, one of which is selected to divide the tree node. Given the number of miRNAs and proteins is large, to reduce computing costs, feature selection, one of the commonly data dimension reduction methods, was applied. It is based on a criterion that selects parts of original features that can best separate different types of samples. According to the feature evaluation strategy used [13], feature selection algorithm can be divided into Filter and Wrapper which are two complementary methods that were combined to characterize the molecular expression levels of normal tissues and tumor tissues. The Filter method is independent of the machine learning algorithm that was subsequently adopted. This method calculates for each feature a statistic that can represent how well a feature has distinguished the sample. On the other hand, the Wrapper method randomly selects a subset of the feature set as a temporary features set for the random forest model, wherein the set with the smallest prediction error and fewer feature numbers serves as the final feature set.

### XGBoost algorithm

XGBoost [14] is similar to Boosting for accurate classification through gradient iterations of weak classifiers. Similar to random forests, each classifier is also a decision tree. Consider this similarity between random forest and XGBoost, to avoid missing important cancer-associated molecules, the results of feature selection of these two algorithms were merged into one set as the final feature set. The construction of the decision tree in the random forest algorithm is independent, however, in the XGBoost algorithm, classifiers are not independent to each other, every latter classifer is optimized based on the classifer result of the previous one. Formally, the mathematical model of XGBoost can be presented as the following formula:

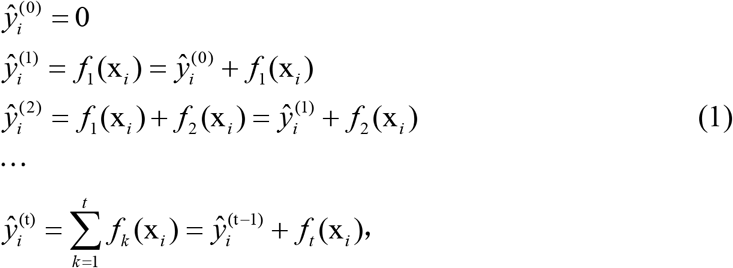

where 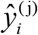 represents the classification result of the first jth classifier. By minimizing the following objective function *Obj*(Θ)^(*t*)^ as follow:

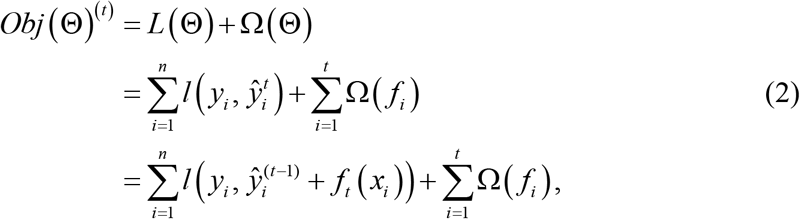

where *L*(Θ) is the loss function to compute error of training set and Ω(Θ) is the regularization term to control complexity of the base classifiers, in the end, the final model can be obtained.

### Maximal information coefficient

Mutual information has been widely used to find non-linear relationships between two variables. Reshef [15] proposed the method of Maximal Information Coefficient (MIC) based on mutual information. The primary advantage of MIC is that a broad correlation analysis can be captured on a sufficient number of statistical samples.*M*(***X***, ***Y***) represents the population feature matrix of ***X***, ***Y***.

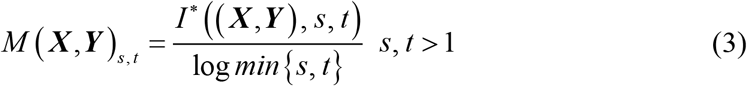

where *I* (***X***, ***Y***) is interactive information of ***X*** and ***Y***, *s*, *t* are the number of divisions on the horizontal and vertical axes, *s · t* < *n^0.6^* (empirical values), and *n* is the number of samples.

In this study, to measure the correlation between any two molecules in the network, the MIC values between any two nodes were calculated. The greater the MIC is, the stronger the correlation is.

### Multilayer network

Multilayer network is denoted by *M*=(*G*,*C*), where *G* = {*G_α_*; *α* ∈{1, 2,…, *m*}} is a set of single layer networks which is denoted as *G_α_* = (*X_α_*, *E_α_*). *X_α_* and *E_α_* is the set of nodes and edges belongs to the layer *G_α_*, respectively. *C* = {*E_αβ_* ⊆ *X_α_ X_β_*;α,β Ω {1, 2,…, *m*}, *α* ≠ *β*} represents the set of edges that connects the nodes in different layers. Elements in *C* are called cross-layer connected edges. Element in *E_α_* is called the intra-layer node connection of *M*. The set of nodes in layer *G_α_* is denoted as: 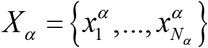, and the adjacency matrix in layer *G_α_* is denoted as:

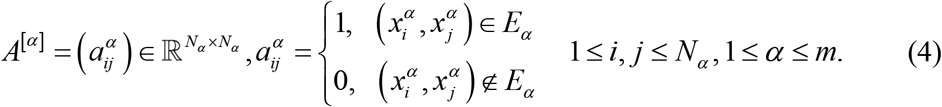

The adjacency matrix in cross-layer *E_αβ_* is denoted as:

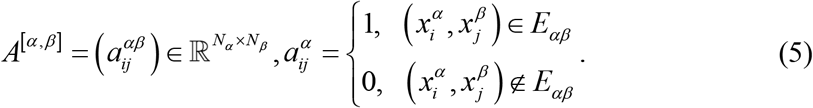

In this study, a single layer network was established between miRNAs, and another single layer network was composed of proteins, which together constituted a two-tier multilayer network. The structure of a multi-layered network can sort out the internal interactions of the same kind of molecules while also taking into account the interactions of different kinds of molecules. Thanks to multilayer structure, in the process of cancer-associated biomarker recognition, the identification of a molecule will no longer be limited to the interaction of the same kind of molecules.

### Betweenness centrality

The betweenness centrality [16] can measure the importance of nodes in the network. If the two network nodes, *v_i_* and *v_j_* are two non-adjacent nodes, the shortest path between them will pass through some nodes. If the certains nodes exists in many of these paths, one can infer that the node is relatively important. The betweenness centrality of node *B_k_* is represented as:

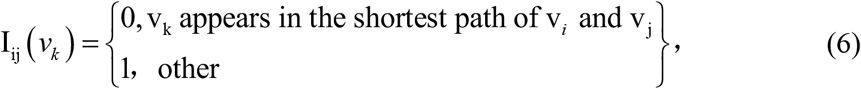

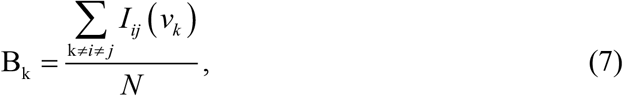

where *N* represents the number of shortest paths. The betweenness centrality reflects the role of the node in the entire network and has a strong practical significance. In different networks, if the betweenness centrality of the same molecule is distinctly different, thereby indicating that this molecule (miRNA or protein) has played a significant role in the breast cancer. In this study we adopted the centrality function of MatLab to calculate the betweenness centrality of the nodes.

### Importance score of nodes

To determine the importance score of nodes in the miRNA layer, 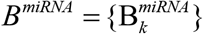 represents the collection of betweenness centrality of nodes in miRNA network. σ(B^miRNA^) represents the standard deviation of the *B^miRNA^* set, and *E*(B^miRNA^) represents the mean of the *B^miRNA^* set. We standardize 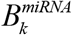 as follows:

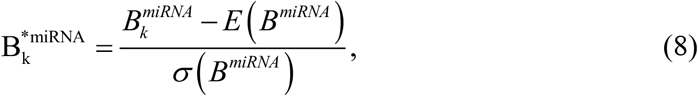

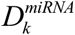 represents the degree of node k in the miRNA network, note that the calculation of degree here only considers cross-layer connected edges:

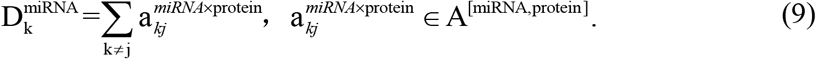

We standardize 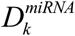 as follows:

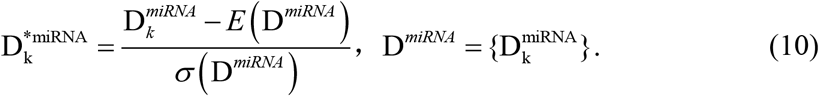

Finally, the importance score of node k in the miRNA network is:

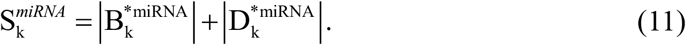

In the same way, calculate the score of the protein molecule as 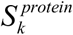.

## Results

### Feature selection

#### XGBoost for feature selection

Because of the imbalance between positive and negative samples of miRNA and protein expression data, up-sampling was used to amplify positive samples. The leave-one-out method was used to train and validate the datasets.

The error rate, logic loss, AMSE of the training and testing datasets (Fig 3 a) gradually decrease in the model, and the AUC (Fig 3 a) gradually increases and stabilizes after 35 iterations. The error rates are 0.0005, 0.005; logical loss values 0.0098, 0.0274; the AUC values close to 1, 1; and the RMSE values 0.0293, 0.0709 in the training and ^1^ testing datasets, respectively. Similarly, the XGBoost algorithm has a high accuracy for ^1^ classification on miRNA expression data. Computation of importance scores (Fig 2b) through the use of XGBoost algorithm suggests that *mir.139*, *mir.21*, *mir.183*, *mir.96*, *mir. 190b* and *mir.6507* are significantly associated with breast cancer.

**Fig 2.**
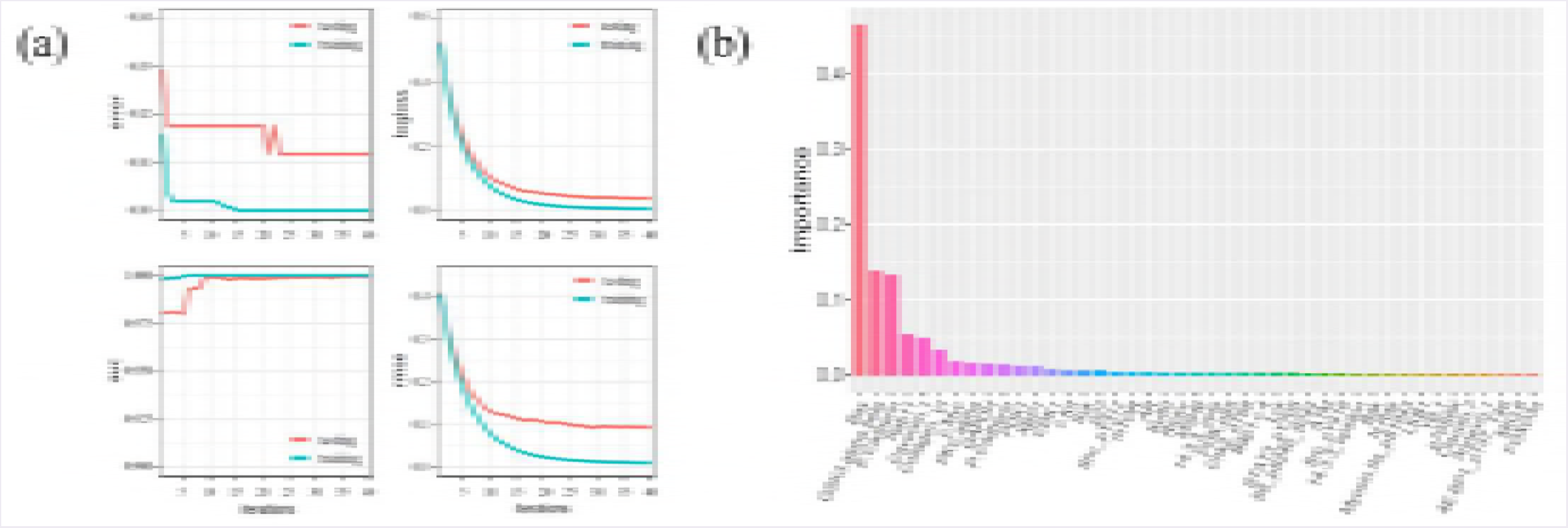
Analysis results of miRNAs based on XGBoost algorithm. **(a)** The trend of error rate, logistic loss, AUC and RMSE in the training and testing of miRNA expression data. Iteration steps ^1^ (x-axis) as well as error rate, logistic loss, AUC and RMSE (y-axis in each of the four panels). Each panel has two lines representing the training set (blue) and test set (red). **(b)** Ranking of important miRNA candidates. miRNA candidates (x-axis) and the importance score of each miRNA candidate (y-axis), as determined through XGBoost algorithm, are shown.

Error rate logic loss and RMSE of the training and testing datasets decrease in the model, whereas the AUC gradually increases then stabilizes after 30 iterations (Fig 3a). The error rates are 0, 0.0118 in the training and testing datasets, respectively; logical loss values are 0.0074, 0.04; AUC values are 1, 0.999; and the RMSE values are 0.0126, 0.0941. Similarly, the XGBoost algorithm is accurate in classifying protein expression data. Importance score as calculated by XGBoost shows that *Bax, GSK3.alpha.beta, E.Cadherin, Rab11, Caveolin.1* and *Collagen_VI* contribute to the high classification accuracy of tumor and normal tissue in breast cancer (Fig 3b).

**Fig 3.**
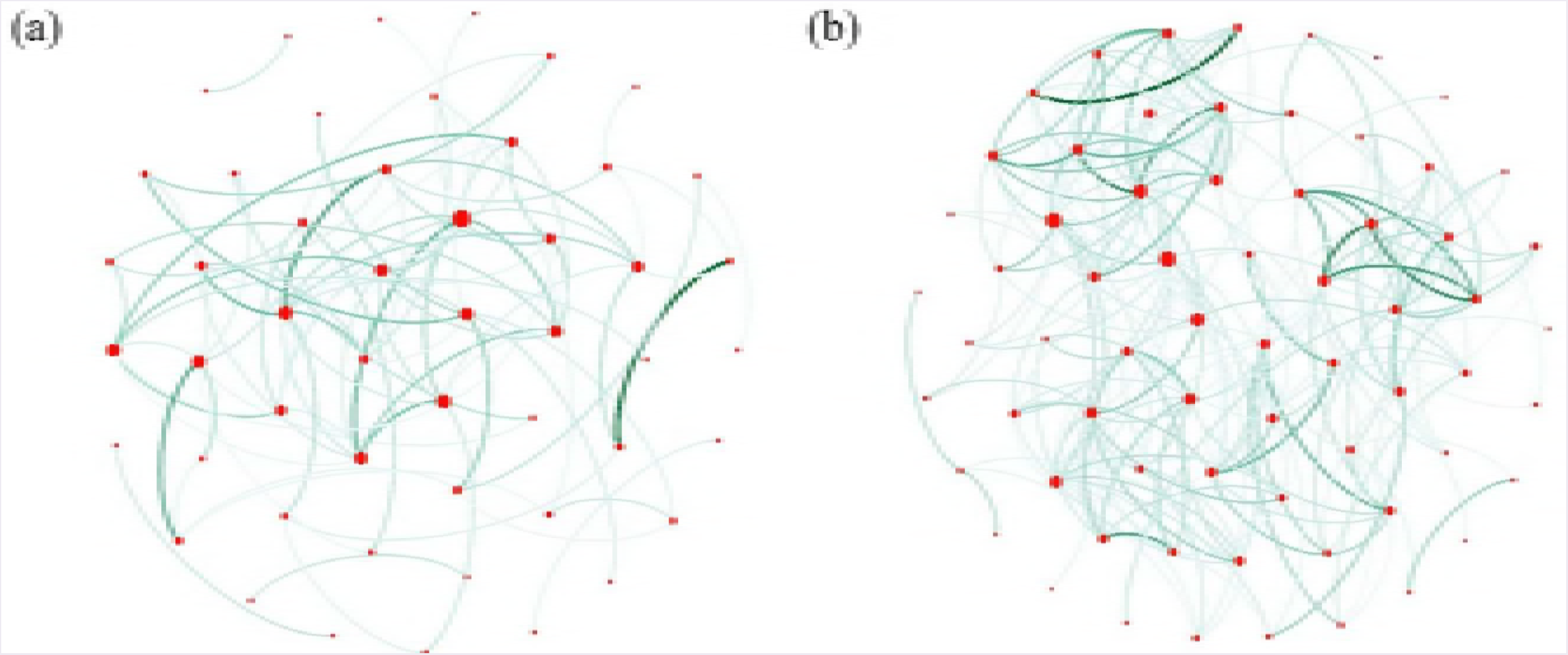
Analysis results of proteins based on XGBoost algorithm. **a** The trend of error rate, logistic loss, auc and rmse in the training and testing of protein expression data. The x-axes represents iteration step and y-axes represents value of error rate, logistic loss, AUC and RMSE, respectively. Every subgraph has two lines represent the training set and test set, respectively. **(b)** The ranking of important variables of protein. The x-axes represents the protein molecules and y-axes represents the importance score of proteins computed by XGBoost algorithm which is different from the score at the end in this study.

XGBoost classification algorithm further shows that some of the classified miRNA (*mir.139* [17], *mir.21* [18], *mir.96* [19], *mir.183* [20]), and protein (*Bax* [21], *GSK3* [22] and *mTOR* [22], *E.cadherin* [23], *Rab11* [24], *caveolin.1* [25]) functions are related to breast cancer.

#### Random forest for feature selection

To estimate the accuracy of the classification, 10-fold cross-validation method was used to assess the classification model (Table 1). When the number of selected miRNAs is 50 in the breast cancer dataset, the cross-validation accuracy rate is 98.50%.

**Table 1.** miRNA classification results by random forest algorithm

| Number of<br>miRNAs | Accuracy<br>coefficient | Kappa<br>coefficient | Accuracy<br>coefficient SD | Kappa<br>coefficient SD |
| --- | --- | --- | --- | --- |
| 10 | 0.9605 | 0.9208 | 0.0314 | 0.06298 |
| 20 | 0.9800 | 0.9600 | 0.02582 | 0.05164 |
| 30 | 0.9755 | 0.9508 | 0.02582 | 0.05164 |
| 50 | 0.9850 | 0.9700 | 0.02415 | 0.04830 |
| 60 | 0.9850 | 0.9700 | 0.02415 | 0.04830 |
| 70 | 0.9800 | 0.9600 | 0.03496 | 0.06992 |
| 80 | 0.9850 | 0.9700 | 0.02415 | 0.04830 |
| 100 | 0.9850 | 0.9700 | 0.02415 | 0.04830 |

The accuracy coefficient measures the correct rate of sample classification, and the Kappa [26] coefficient is used for checking consistency and could also measure the effect of classification accuracy. As accuracy and Kappa coefficients increase, their standard deviations decrease (Table 1).

In this study, four cancer-associated miRNAs were screened by XGBoost algorithm, and three of them, namely *mir.21*, *mir.96*, and *mir.183*, were screened out by random forests. A comparison of the two miRNA datasets indicated that 28% of the feature selections are consistent. Similar to the analysis miRNA datasets, a 10-fold cross-validation method was used to assess the classification model to obtain protein classification (Table 2).

**Table 2.** Protein classification results by random forest algorithm

| Number of Proteins | Accuracy | Kappa | Accuracy SD | Kappa SD |
| --- | --- | --- | --- | --- |
| 10 | 0.9476 | 0.8952 | 0.09024 | 0.1770 |
| 20 | 0.9342 | 0.8691 | 0.09068 | 0.1789 |
| 30 | 0.9342 | 0.8691 | 0.09068 | 0.1789 |
| 50 | 0.9231 | 0.8448 | 0.10284 | 0.2081 |
| 60 | 0.9231 | 0.8448 | 0.10284 | 0.2081 |
| 70 | 0.9231 | 0.8448 | 0.10284 | 0.2081 |
| 78 | 0.9231 | 0.8448 | 0.10284 | 0.2081 |

For breast cancer datasets, when the number selected proteins is 10, the cross-validation accuracy is 94.76%. In the 10 selected proteins, *Bax* [21], *GSK3* [22], *E.cadherin* [23], *caveolin.1* [25], *PI3K* [27], *Collagen* [28], *XBP1* [29], *syk* [30] were found to be significantly associated with breast cancer.

### Summary of feature selection

After obtaining two miRNA candidate sets and two protein candidate sets selected by two algorithms, the union of the two miRNA sets was taken as the final miRNA candidate set, in the same manner, the final proteins candidate set was obtained. The number of selected miRNA sets is 86, and the number of selected protein sets is 30.

### Calculate MIC and threshold setting

As the MIC increases, the number of nodes and edges decreases. If the selected MIC threshold is so small that the number of nodes and edges in both network becomes too large, identification of nodes that have significant differences becomes more difficult. If the selected MIC threshold is so large that the network becomes too sparse, many connections are missed, which is not conducive to analyze the relationship between the nodes. MIC was calculated between any two candidates in the miRNA and protein datasets obtained through feature selection, and the threshold was set to 0.2, 0.35, and 0.5.

Under the MIC threshold of 0.5, miRNA network of cancer tissue and normal tissue was plotted (Fig 4). The cancer network (Fig 4a) is sparser than the normal network (Fig 4b). This finding indicates that the interaction of miRNA networks differs significantly by cell type and supports the use of complex networks for breast cancer analysis. Similarly, under the MIC threshold of 0.5, protein network of cancer tissue and normal tissue was also plotted.

**Fig 4.**
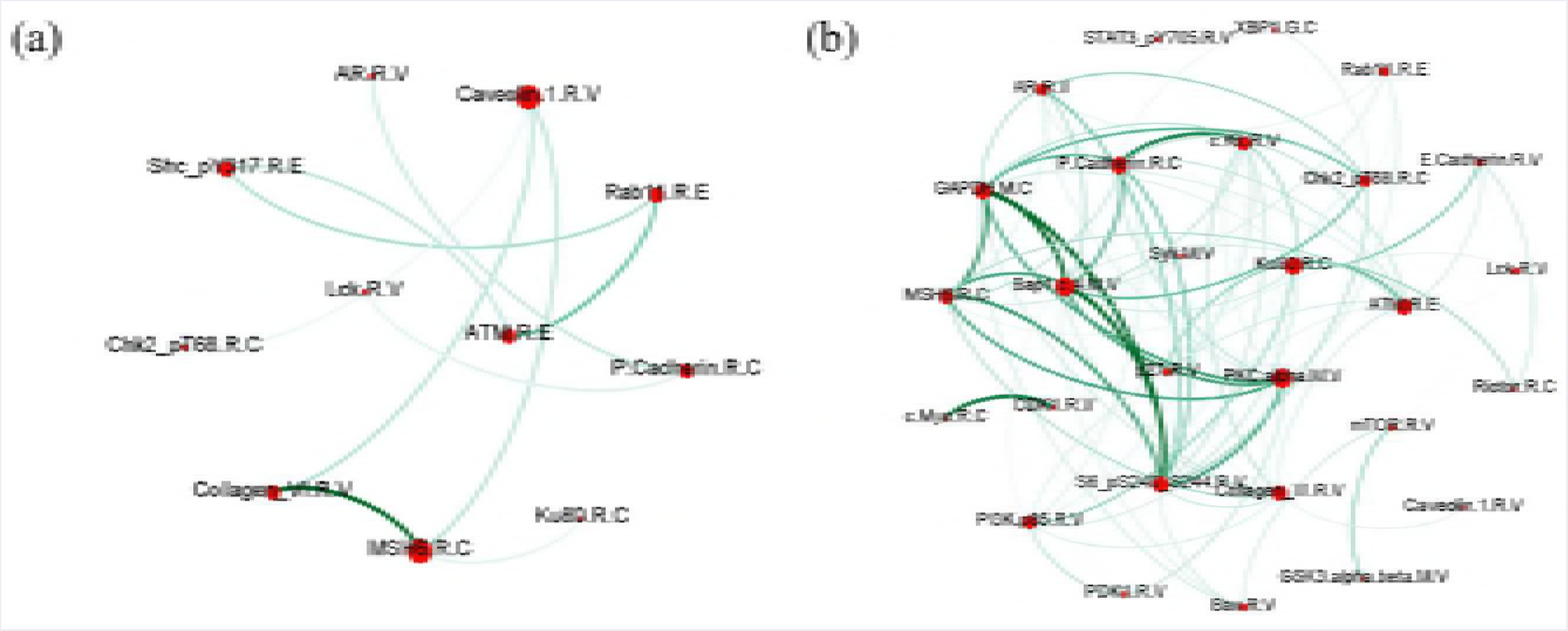
miRNA network of cancer tissue and normal tissue. **(a)** miRNA network of cancer tissue containing candidates with MIC greater than 0.5; **(b)** miRNA network of normal tissue containing MIC greater than 0.5. The size of the node represents node degree which is the number of connections it has to other nodes and the color datkness of the edge represents the size of the MIC value.

The protein network also shows the same characteristics as miRNAs—namely, that the network of cancerous tissue is much sparser than that of normal tissue (Fig 5). Because the miRNA network of cancer cells has a small number of nodes at an MIC threshold of 0.5 and may miss some important proteins, we decided not to use this MIC threshold to construct a complex network.

**Fig 5.**
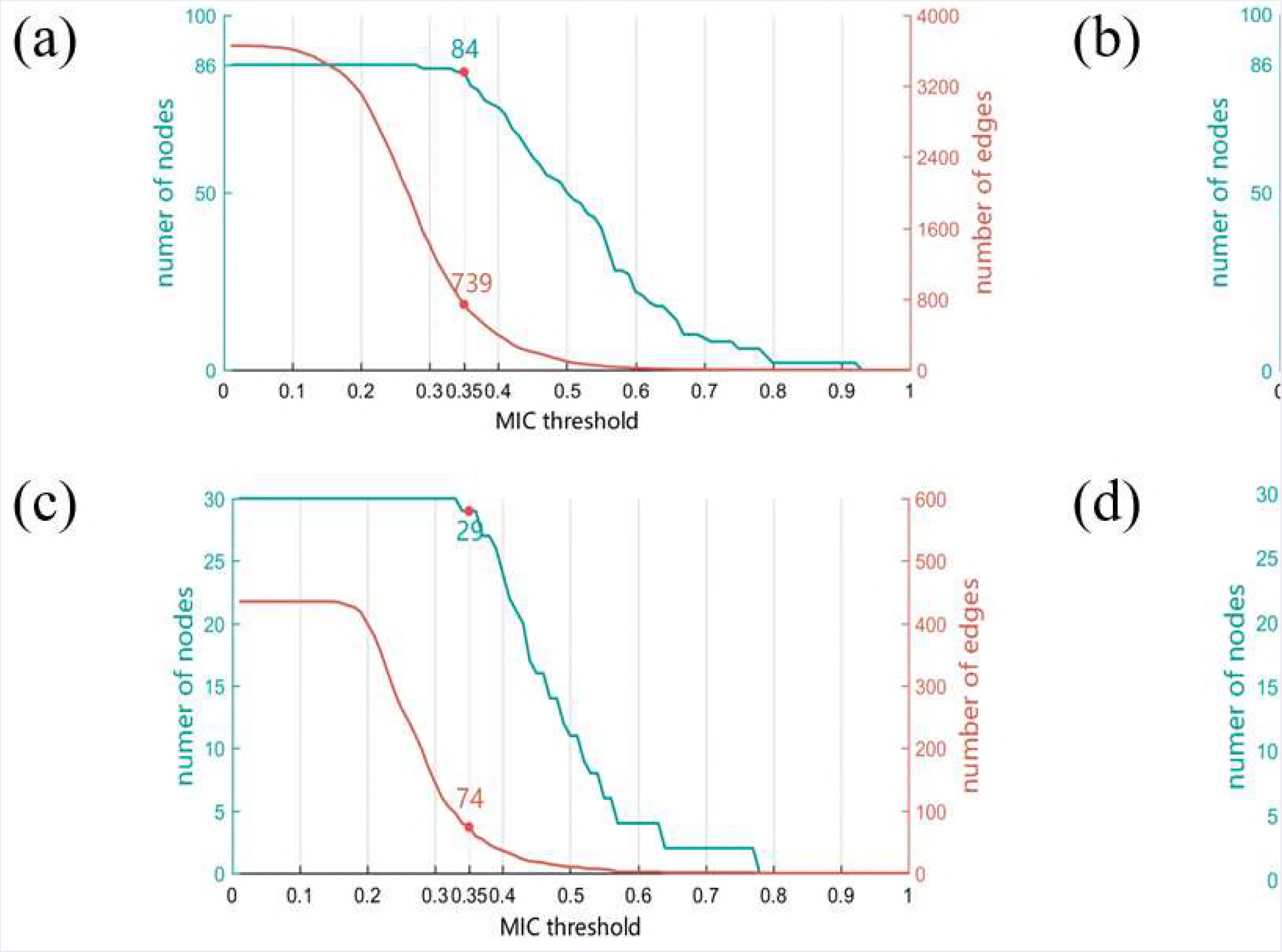
Protein network of cancer tissue and normal tissue. **(a)** Protein network of cancer tissue containing candidates with MIC greater than 0.5; **(b)** Protein network of normal tissue containing candidates with MIC greater than 0.5. The size of the node represents the size of the degree, and the color depth of the edge represents the size of the MIC value.

Figure analysis was applied to determine which MIC threshold should be adopted. While the number of nodes varies inappreciably when MIC threshold is set to 0.35, the number of connections between nodes is significantly reduced, suggesting that these complex networks are distinct.

Several principles were considered when selecting a threshold. Firstly, the MIC threshold selected must not lead to the loss of too many nodes. For instance, in the analytical process described, less than 5% of nodes are lost. Secondly, the number of edges of the network could not be too small, and the number of edges of the two networks must be significantly different. Number of edges in the miRNA network of normal tissue (Fig 6b) is about 1.59 times that in the miRNA network of cancer tissue (Fig 6a).

**Fig 6.**
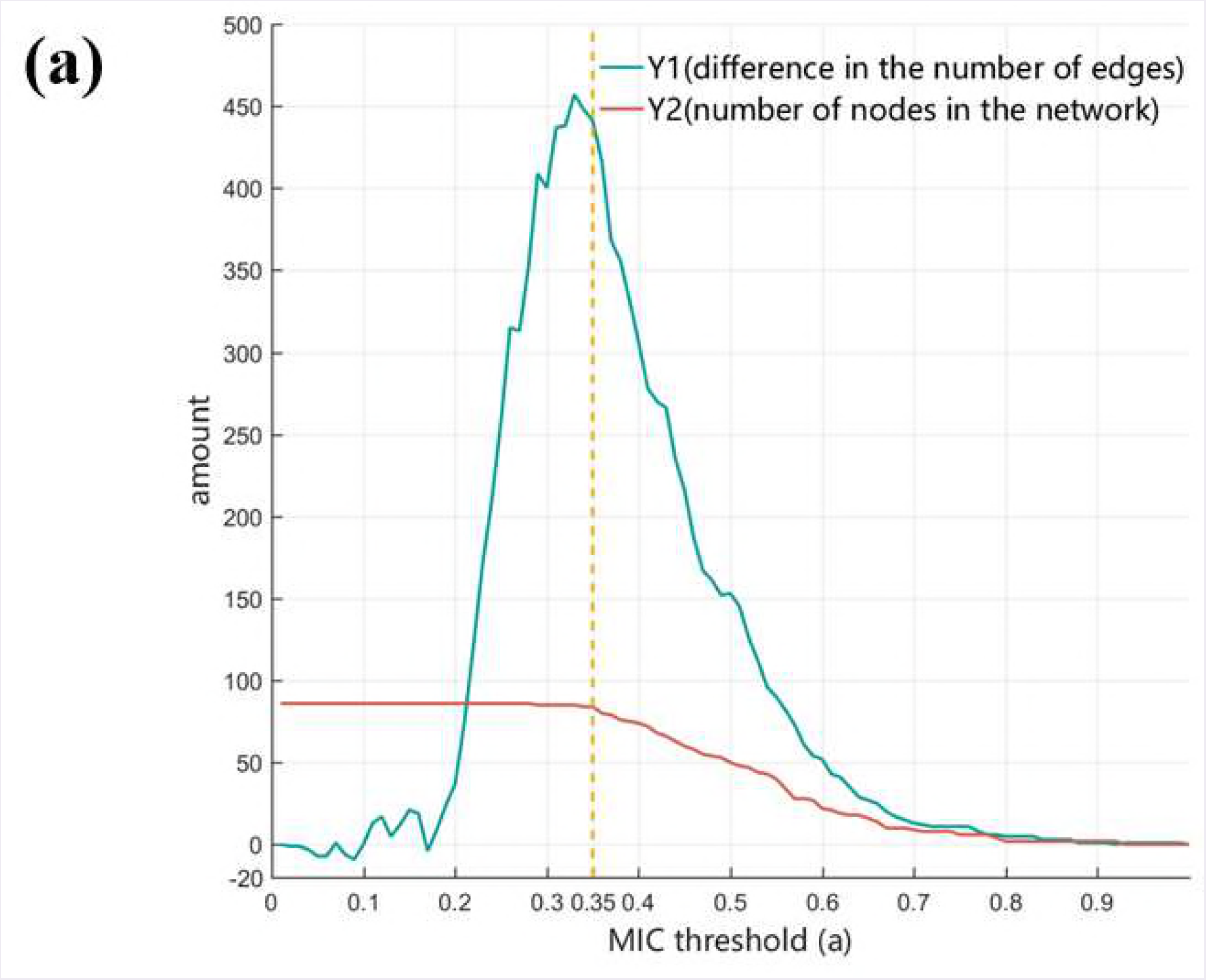
Relationship between MIC threshold and the number of nodes and edges. Analyses of miRNA networks of cancer tissue (a) and normal tissue (b), as well as protein networks of cancer tissue (c) and normal tissue (d) are shown. The x-axes represent threshold of MIC, y-axis to the left of each figure represents number of nodes and y-axis to the right represents number of edges in the network. Each figure shows the number of nodes or edges corresponding to a threshold of 0.35.

Through observing the difference in structure of network under different thresholds (Fig 7), it was be found that when the MIC threshold is 0.35, the decrease in the number of network nodes is not obvious, effectively fulfilling the first principle of threshold selection. Difference in the number of edges between the two networks is also kept at a relatively high level; that is, this MIC threshold could effectively differentiate the two networks, which is in accordance with the second principle of threshold selection. Therefore, we selected 0.35 as the MIC threshold for analysis.

**Fig 7.**
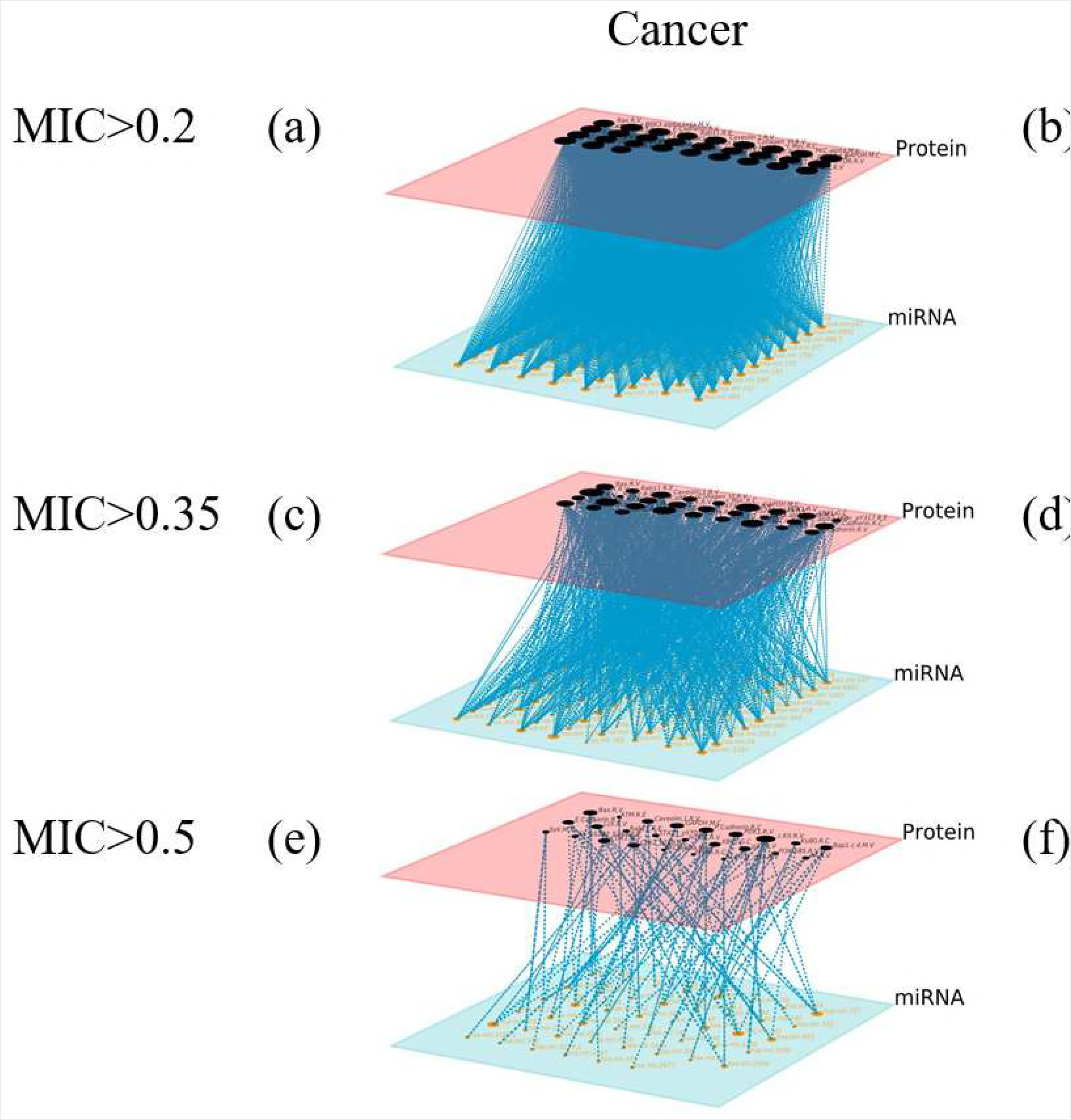
Relationship between MIC threshold and structure differences within the (a) miRNA network and (b) protein network. Curve Y1, which is obtained by subtracting the number of edges of the normal tissue and the cancer tissue network, represents the structural difference between the two networks. Curve Y2, which is obtained by selecting the smaller values of the number of nodes of the normal tissue and the cancer tissue network, represents the richness of the nodes of both networks. The x-axis of each figure denotes threshold of MIC and the y-axis indicates the value corresponding to Y1 and Y2.

### Multilayer network

Multilayer networks were generated after calculating MIC between nodes and setting MIC thresholds to 0.2, 0.35, or 0.5 (Fig 8).

**Fig 8.**
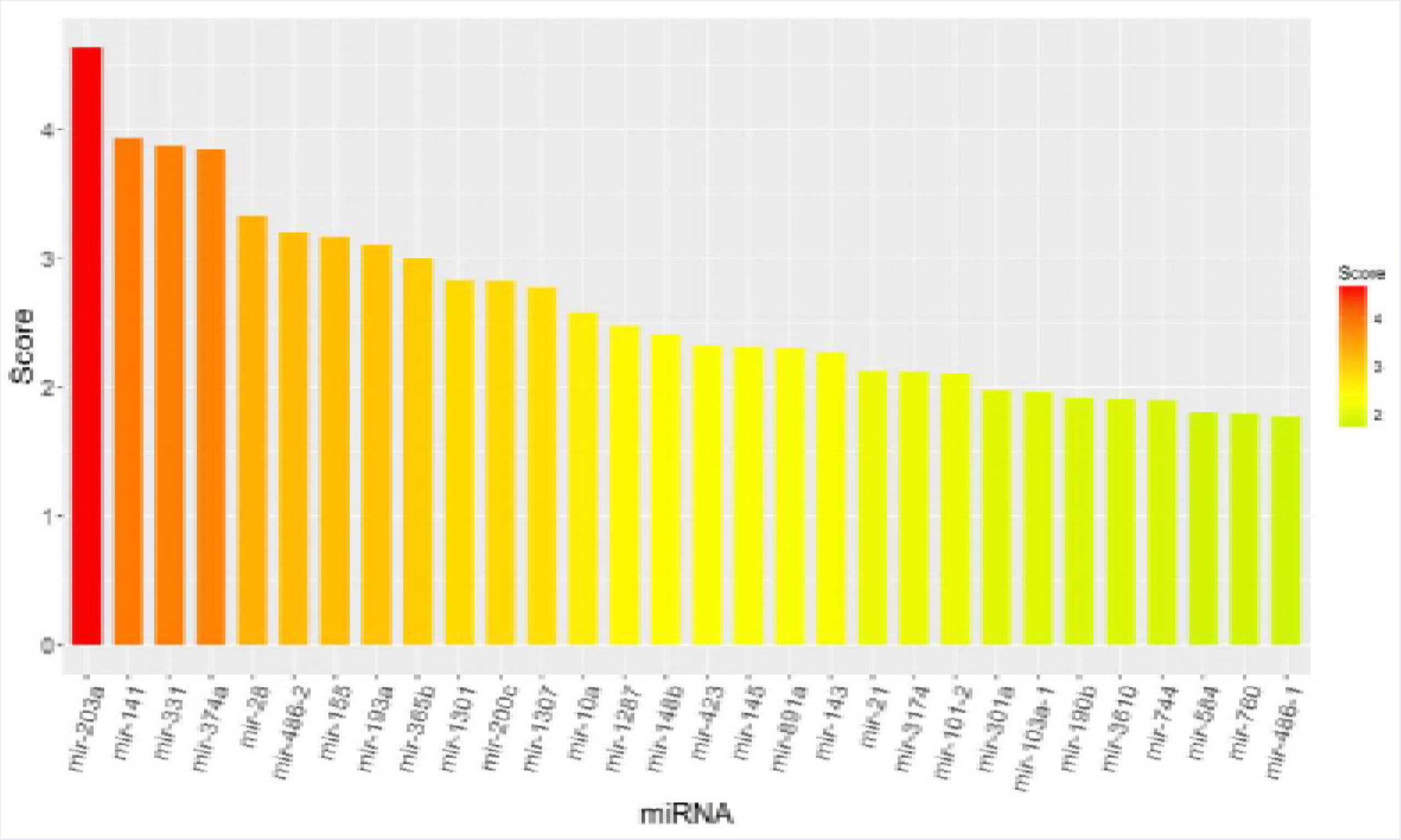
The multilayer network of cancer tissue and normal tissue. The multilayer networks were constructed for cancer tissue (a, c, e) and normal tissue (b, d, f) with MIC greater than 0.2 (a & b), MIC greater than 0.35 (c & d), and MIC greater than 0.5 (e & f). The red layer is the protein layer and the blue layer is the miRNA layer.

A multilayer network with a threshold of 0.2 had more edges than other multilayer network with higher threshold, which causes the impact of key connections in the network become smaller. Hence, under the threshold of 0.2, two types of cells is difficult to be distinguished well with this threshold. When the threshold is 0.5, there are obvious differences between the two multi-layer networks, but the number of edges is sparse and some important relationships may be mistakenly omitted. These problems are averted when the threshold of 0.35 is used, which affirms the use of this threshold.

### Node ranking

To better understand the details of the networks, the nodes representing different candidates were ranked according to betweenness centrality and node degree (Figs 9 and 10).

**Fig 9.**
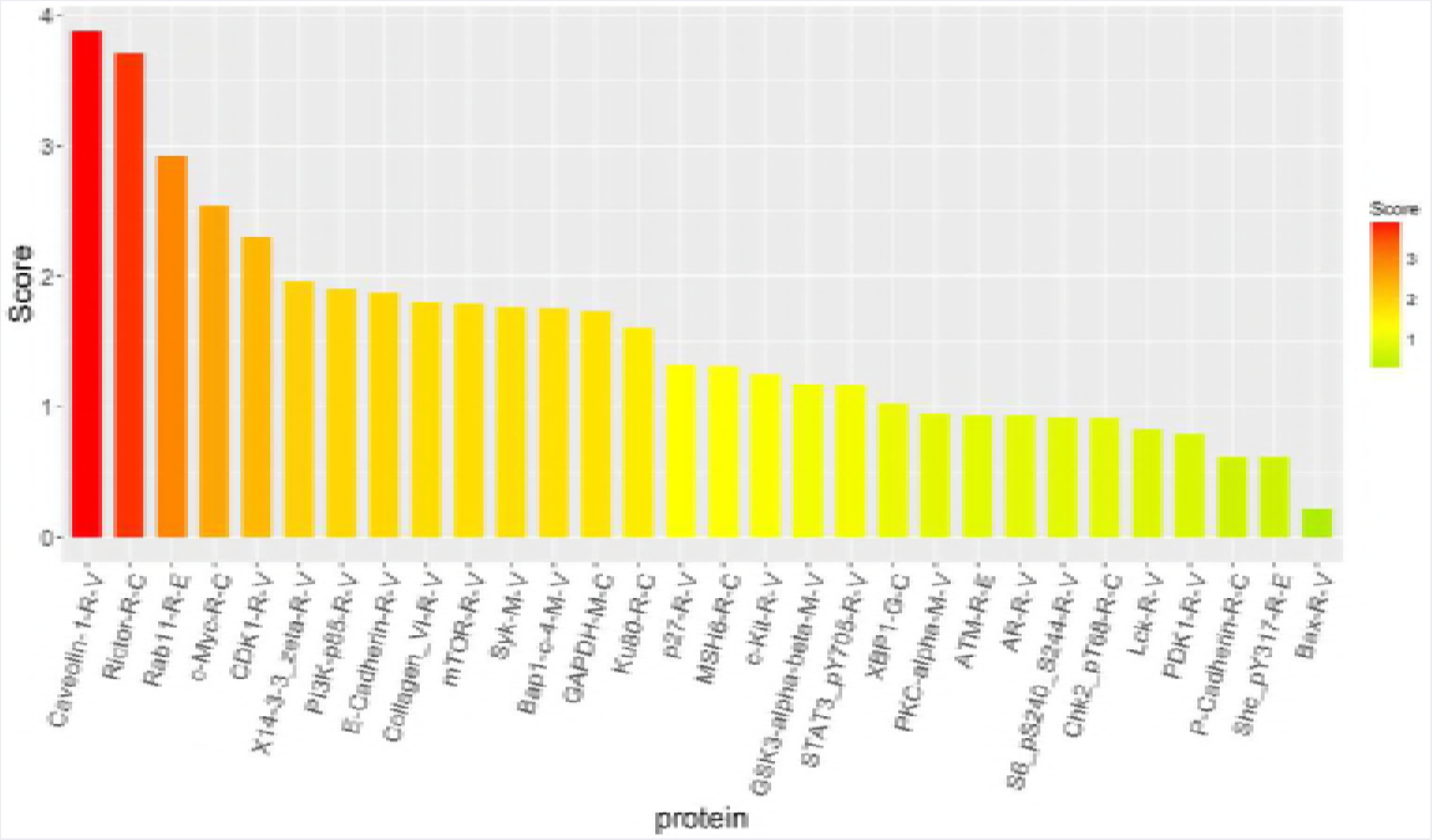
Score ranking of miRNAs. The horizontal axis represents each molecule (top 30 scores), and the vertical axis represents the magnitude of the importance score. The x-axis represents miRNA molecule, and y-axis represents score of miRNAs.

**Fig 10.**
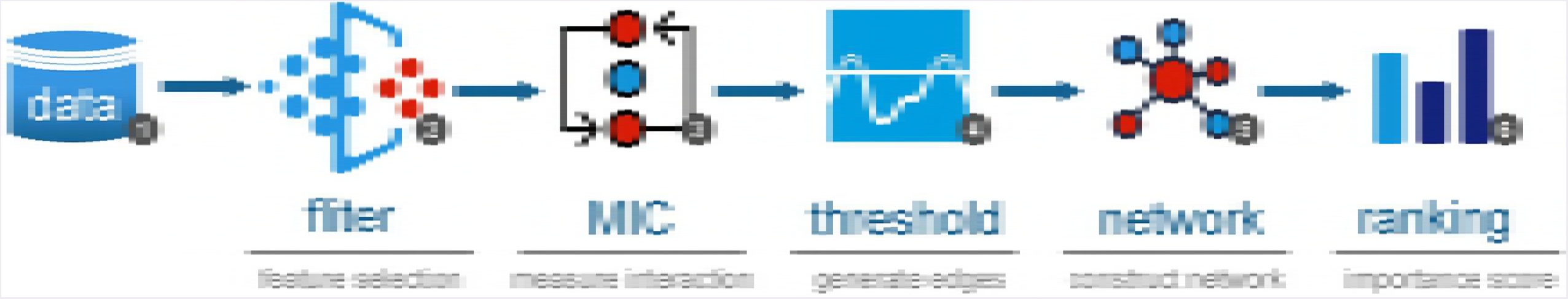
Score ranking of Protein. The horizontal axis represents each molecule, and the vertical axis represents the magnitude of the importance score. The x-axes represents miRNA molecule and y-axes represents score of proteins.

Among the top 15 miRNAs, the relationships of 11 miRNAs with breast cancer in previously published studies were confirmed (Table 3).

**Table 3.** Cancer-related miRNAs

| miRNA Candidate | Description |
| --- | --- |
| *mir-203a* [31] | Reconstitution of *Runx2* in *MDA-MB-231-luc* cells delivered with *miR-203* reverses the inhibitory effect of the miRNAs on tumor growth and metastasis. |
| *mir-141* [32] | *miR-141* has distinct profiles in *EGF-dependent* breast cancer cell invasion, proliferation, and cell cycle progression. |
| *mir-374a* [33] | The *Wnt/β-catenin* signaling is hyperactivated in metastatic breast cancer cells that express *miR-374a*. |
| *mir-28* [34] | In breast cancer cells, *miR-28* regulates *Nrf2* expression at the posttranscriptional level by binding to the 3'UTR of *Nrf2* mRNA and resulting in *Nrf2* mRNA degradation. |
| *mir-155* [35] | *miR-155* expression is upregulated in breast cancer cells, which reduces the levels of *RAD51* and affects the cellular response to ionizing radiation. |
| *mir-193a* [36] | *miR-193a* expression is downregulated in breast cancer cell lines and tissues when compared with the adjacent non-tumor tissues. |
| *mir-365b* [37] | *miR-365* expression levels are significantly higher in breast cancer tissues when compared with adjacent non-tumor tissues. |
| *mir-1301* [38] | *miR-1301* is overexpressed in breast cancer tissues and cell lines and cell tissues, whereas downregulation of *miR-1301* inhibits the proliferation of breast cancer cells in vitro. |
| *mir-200c* [39] | For the claudin-low breast cancer, *miR-200c* has therapeutic effects in an in vivo model. |
| *mir-10a* [40] | The median expression levels of *miR-10b* in tumor tissue when compared with adjacent non-tumor tissue are significantly higher in relapsed patients than in relapse-free patients. |
| *mir-148b* [41] | *miR148b* is a major coordinator of breast cancer progression in a relapse-associated microRNA signature. |

Among the selected proteins, we were able to confirm the relationships of the top 10 proteins with breast cancer using published literature (Table 4).

**Table 4.** Cancer-related Proteins

| Protein Candidate | Description |
| --- | --- |
| *E-Cadherin-R-V* [23] | *E-cadherin* is regulated epigenetically via methylation of the promoter in most intraductal breast carcinomas. |
| *Caveolin-1-R-V* [25] | *Caveolin-1* expression is significantly decreased in breast cancer-associated fibroblasts compared to normal fibroblasts and is associated with increased invasion-promoting capacity. |
| *PI3K-p85-R-V* [27] | The inhibition of *PI3K* promotes ER activity, as manifested by increases in ER binding to target promoters and ER target gene expression. |
| *Collagen_VI-R-V* [28] | *Collagen VI* are upregulated in breast cancer, generating a microenvironment that promotes tumour progression and metastasis. |
| *Rictor-R-C* [42] | *Rictor* expression is upregulated significantly as compared with nonmalignant tissues in invasive breast cancer specimens. |
| *Rab11-R-E* [43] | Rab coupling protein (*FIP1C*), an effector of the *Rab11* GTPases, is amplified and overexpressed in 10% to 25% of primary breast cancers. |
| *c-Myc-R-C* [44] | In the *AhR/HDAC6/c-Myc* signaling pathway, phthalates induce proliferation and invasiveness of estrogen receptor-negative breast cancer. |
| *CDK1-R-V* [45] | Combined inhibition of *Cdk1* and *PARP* in BRCA–wild-type cancer cells (breast cancer–associated cell) results in reduced colony |

## Discussion

The multilayer network analysis proposed helps identify miRNAs and proteins that could be associated with breast cancer. While biomarkers were previously selected using machine learning, this study is novel in that a combination of machine learning and multilayer network methods was used.

This combinatorial approach to identifying cancer biomarkers could prevent missing critical miRNA or protein candidates and ensure a more robust analysis. For example, the final ranking of nodes generated from multilayer network method in combination with machine learning differs from that using machine learning alone. This finding suggests that the combinatorial effect of multilayer network analysis and machine learning yields more comprehensive information. It also shows that the multilayer network analysis method could facilitate the discovery of novel molecular candidates.

Although published work has described miRNA or protein networks of expression profiles separately, interrelationships between two networks have not been thoroughly investigated. To address this knowledge gap, the interrelationship between miRNA and protein networks was studied through the MIC. The most suitable MIC threshold was determined by analyzing the interrelationship between the MIC threshold and the number of nodes and edges of the network. By using the most optimized MIC threshold to construct the multilayer networks, miRNAs and proteins associated with breast cancer were identified. Although the top-ranked candidates for protein biomarkers were previously identified, the combinatorial approach proposed reveals potentially novel miRNAs associated with breast cancer, such as mir-331, mir-486-2, mir-1307 and mir-1287. Roles of these miRNA candidates in breast cancer will be confirmed through molecular means.

A minor drawback associated with the multilayer network analysis is that the optimized threshold value must be determined through analysis of the distribution map (Figs 6 and 7) measuring the number of edges and nodes in network under different thresholds. The approximate range can be selected but the optimal value cannot be obtained automatically. This shortcoming could be overcome by establishing algorithms that facilitate selection of optimized threshold values.

Because single network analysis provides limited information, the proposed combinatorial approach will allow for a deeper understanding of multiple networks and signaling pathway in cancer. The regulatory architecture of miRNA and protein in breast cancer patients analyzed in multiple network-wide will potentially enable novel cancer biomarker discovery.

## Supporting information

### S1 File. miRNA expression data of breast cancer

The first column of the data is the name of miRNAs, and first row of the data is the code of the corresponding cases of samples in TCGA, where the fields ending with ‘-11’ is normal samples and the rest are cancer tissues.

### S2 File. Protein expression data of breast cancer

The first column of the data represent cell type of the miRNAs, where ‘0’ means cancer samples and ‘1’ means normal tissues. The second column of the data is the code of the corresponding cases of samples in TCGA, and first row of the data is the name of proteins.

## References

[1] Ferlay J, Shin HR, Bray F, Forman D, Mathers C, Parkin DM. Estimates of worldwide burden of cancer in 2008: GLOBOCAN 2008. International journal of cancer. 2010 Dec 15;127(12):2893–917.

[2] Lin S, Gregory RI. MicroRNA biogenesis pathways in cancer. Nature reviews cancer. 2015 Jun;15(6):321.

[3] Acunzo M, Romano G; Wernicke D, Croce CM. MicroRNA and cancer-a brief overview. Advances in biological regulation. 2015 Jan 1;57:1–9.

[4] Lee YS, Dutta A. MicroRNAs in cancer. Annual Review of Pathological Mechanical Disease. 2009 Feb 28;4:199–227.

[5] Piletič K, Kunej T. MicroRNA epigenetic signatures in human disease. Archives of toxicology. 2016 Oct 1;90(10):2405–19.

[6] Shruthi BS, Palani Vinodhkumar S. Proteomics: A new perspective for cancer. Advanced biomedical research. 2016;5.

[7] Gam LH. Breast cancer and protein biomarkers. World journal of experimental medicine. 2012 Oct 20;2(5):86.

[8] Obermeyer Z, Emanuel EJ. Predicting the future—big data, machine learning, and clinical medicine. The New England journal of medicine. 2016 Sep 29;375(13):1216.

[9] Leung MK, Delong A, Alipanahi B, Frey BJ. Machine learning in genomic medicine: a review of computational problems and data sets. Proceedings of the IEEE. 2016 Jan;104(1):176–97.

[10] Wang J, Yang X, Cai H, Tan W, Jin C, Li L. Discrimination of breast cancer with microcalcifications on mammography by deep learning. Scientific reports. 2016 Jun 7;6:27327.

[11] Wu B, Abbott T, Fishman D, McMurray W, Mor G, Stone K, Ward D, Williams K, Zhao H. Comparison of statistical methods for classification of ovarian cancer using mass spectrometry data. Bioinformatics. 2003 Sep 1;19(13):1636–43.

[12] Biau G Scornet E. A random forest guided tour. Test. 2016 Jun 1;25(2):197–227.

[13] Inza I, Larrañaga P, Blanco R, Cerrolaza AJ. Filter versus wrapper gene selection approaches in DNA microarray domains. Artificial intelligence in medicine. 2004 Jun 1;31(2):91–103.

[14] Chen T, Guestrin C. Xgboost: A scalable tree boosting system. InProceedings of the 22nd acm sigkdd international conference on knowledge discovery and data mining 2016 Aug 13 (pp. 785–794). ACM.

[15] Reshef DN, Reshef YA, Finucane HK, Grossman SR, McVean Q Turnbaugh PJ, Lander ES, Mitzenmacher M, Sabeti PC. Detecting novel associations in large data sets. science. 2011 Dec 16;334(6062):1518–24.

[16] Borgatti SP. Centrality and network flow. Social networks. 2005 Jan 1;27(1):55–71.

[17] Krishnan K, Steptoe AL, Martin HC, Pattabiraman DR, Nones K, Waddell N, Mariasegaram M, Simpson PT, Lakhani SR, Vlassov A, Grimmond SM. miR-139-5p is a regulator of metastatic pathways in breast cancer. Rna. 2013 Dec 1;19(12):1767–80.

[18] Yan LX, Huang XF, Shao Q, Huang MY, Deng L, Wu QL, Zeng YX, Shao JY. MicroRNA miR-21 overexpression in human breast cancer is associated with advanced clinical stage, lymph node metastasis and patient poor prognosis. Rna. 2008 Nov 1;14(11):2348–60.

[19] Hong Y, Liang H, Wang Y, Zhang W, Zhou Y, Yu M, Cui S, Liu M, Wang N, Ye C, Zhao C. miR-96 promotes cell proliferation, migration and invasion by targeting PTPN9 in breast cancer. Scientific reports. 2016 Nov 18;6:37421.

[20] Macedo T, Silva-Oliveira RJ, Silva VA, Vidal DO, Evangelista AF, Marques M. sOverexpression of mir-183 and mir-494 promotes proliferation and migration in human breast cancer cell lines. Oncology letters. 2017 Jul 1;14(1):1054–60.

[21] Kholoussi NM, El-Nabi SE, Esmaiel NN, Abd El-Bary NM, El-Kased AF. Evaluation of Bax and Bak gene mutations and expression in breast cancer. BioMed research international.2014;2014.

[22] Azoulay-Alfaguter I, Elya R, Avrahami L, Katz A, Eldar-Finkelman H. Combined regulation of mTORC1 and lysosomal acidification by GSK-3 suppresses autophagy and contributes to cancer cell growth. Oncogene. 2015 Aug;34(35):4613.

[23] Chao YL, Shepard CR, Wells A. Breast carcinoma cells re-express E-cadherin during mesenchymal to epithelial reverting transition. Molecular cancer. 2010 Dec;9(1):179.

[24] Boulay PL, Mitchell L, Turpin J, Huot-Marchand JÉ, Lavoie C, Sanguin-Gendreau V, Jones L, Mitra S, Livingstone JM, Campbell S, Hallett M. Rab11-FIP1C is a critical negative regulator in ErbB2-mediated mammary tumor progression. Cancer research. 2016 May 1;76(9):2662–74.

[25] Simpkins SA, Hanby AM, Holliday DL, Speirs V. Clinical and functional significance of loss of caveolin-1 expression in breast cancer-associated fibroblasts. The Journal of pathology. 2012 Aug 1;227(4):490–8.

[26] Viera AJ, Garrett JM. Understanding interobserver agreement: the kappa statistic. Fam Med. 2005 May 1;37(5):360–3.

[27] Bosch A, Li Z, Bergamaschi A, Ellis H, Toska E, Prat A, Tao JJ, Spratt DE, Viola-Villegas NT, Castel P, Minuesa G. PI3K inhibition results in enhanced estrogen receptor function and dependence in hormone receptor-positive breast cancer. Science translational medicine. 2015 Apr 15;7(283):283ra51-.

[28] Karousou E, DAngelo ML, Kouvidi K, Vigetti D, Viola M, Nikitovic D, De Luca G Passi A. Collagen VI and hyaluronan: the common role in breast cancer. BioMed research international. 2014;2014.

[29] Chen X, Iliopoulos D, Zhang Q, Tang Q, Greenblatt MB, Hatziapostolou M, Lim E, Tam WL, Ni M, Chen Y, Mai J. XBP1 promotes triple-negative breast cancer by controlling the HIFla pathway. Nature. 2014 Apr;508(7494):103.

[30] Hardy SD, Geahlen RL. Investigating the role of Syk in TGF-β induced P-bodies and breast cancer metastasis.

[31] Taipaleenmäki H, Browne G; Akech J, Zustin J, Van Wijnen AJ, Stein JL, Hesse E, Stein GS, Lian JB. Targeting of Runx2 by miR-135 and miR-203 impairs progression of breast cancer and metastatic bone disease. Cancer research. 2015 Apr 1;75(7):1433–44.

[32] Uhlmann S, Zhang JD, Schwäger A, Mannsperger H, Riazalhosseini Y, Burmester S, Ward A, Korf U, Wiemann S, Sahin Ö. miR-200bc/429 cluster targets PLCyl and differentially regulates proliferation and EGF-driven invasion than miR-200a/141 in breast cancer. Oncogene. 2010 Jul;29(30):4297.

[33] Cai J, Guan H, Fang L, Yang Y, Zhu X, Yuan J, Wu J, Li M. MicroRNA-374a activates Wnt/β-catenin signaling to promote breast cancer metastasis. The Journal of clinical investigation. 2013 Jan 16;123(2).

[34] Yang M, Yao Y, Eades G; Zhang Y, Zhou Q. MiR-28 regulates Nrf2 expression through a Keap1-independent mechanism. Breast cancer research and treatment. 2011 Oct 1; 129(3):983–91.

[35] Gasparini P, Lovat F, Fassan M, Casadei L, Cascione L, Jacob NK, Carasi S, Palmieri D, Costinean S, Shapiro CL, Huebner K. Protective role of miR-155 in breast cancer through RAD51 targeting impairs homologous recombination after irradiation. Proceedings of the National Academy of Sciences. 2014 Mar 25;111(12):4536–41.

[36] Xie F, Hosany S, Zhong S, Jiang Y, Zhang F, Lin L, Wang X, Gao S, Hu X. MicroRNA-193a inhibits breast cancer proliferation and metastasis by downregulating WT1. PloS one. 2017 Oct 10;12(10):e0185565.

[37] Li M, Liu L, Zang W, Wang Y, Du Y, Chen X, Li P, Li J, Zhao G. miR-365 overexpression promotes cell proliferation and invasion by targeting ADAMTS-1 in breast cancer. International journal of oncology. 2015 Jul 1;47(1):296–302.

[38] Lin WH, Li J, Zhang B, Liu LS, Zou Y, Tan JF, Li HP. MicroRNA-1301 induces cell proliferation by downregulating ICAT expression in breast cancer. Biomedicine & Pharmacotherapy. 2016 Oct 1;83:177–85.

[39] Knezevic J, Pfefferle AD, Petrovic I, Greene SB, Perou CM, Rosen JM. Expression of miR-200c in claudin-low breast cancer alters stem cell functionality, enhances chemosensitivity and reduces metastatic potential. Oncogene. 2015 Dec;34(49):5997.

[40] Chang CH, Fan TC, Yu JC, Liao GS, Lin YC, Shih AC, Li WH, Yu AL. The prognostic significance of RUNX2 and miR-10a/10b and their inter-relationship in breast cancer. Journal of translational medicine. 2014 Dec;12(1):257.

[41] Zhang JG; Shi Y, Hong DF, Song M, Huang D, Wang CY, Zhao G. MiR-148b suppresses cell proliferation and invasion in hepatocellular carcinoma by targeting WNT1/β-catenin pathway. Scientific reports. 2015 Jan 28;5:8087.

[42] Joly MM, Hicks DJ, Jones B, Sanchez V, Estrada MV, Young C, Williams M, Rexer BN, Sarbassov DD, Muller WJ, Brantley-Sieders D. Rictor/mTORC2 drives progression and therapeutic resistance of HER2-amplified breast cancers. Cancer research. 2016 Aug 15;76(16):4752–64.

[43] Boulay PL, Mitchell L, Turpin J, Huot-Marchand JÉ, Lavoie C, Sanguin-Gendreau V, Jones L, Mitra S, Livingstone JM, Campbell S, Hallett M. Rab11-FIP1C is a critical negative regulator in ErbB2-mediated mammary tumor progression. Cancer research. 2016 May 1;76(9):2662–74.

[44] Hsieh TH, Tsai CF, Hsu CY, Kuo PL, Lee JN, Chai CY, Wang SC, Tsai EM. Phthalates induce proliferation and invasiveness of estrogen receptor-negative breast cancer through the AhR/HDAC6/c-Myc signaling pathway. The FASEB Journal. 2012 Feb 1;26(2):778–87.

[45] Johnson N, Li YC, Walton ZE, Cheng KA, Li D, Rodig SJ, Moreau LA, Unitt C, Bronson RT, Thomas HD, Newell DR. Compromised CDK1 activity sensitizes BRCA-proficient cancers to PARP inhibition. Nature medicine. 2011 Jul;17(7):875.

[46] Lu J, Guo H, Treekitkarnmongkol W, Li P, Zhang J, Shi B, Ling C, Zhou X, Chen T, Chiao PJ, Feng X. 14-3-3Z cooperates with ErbB2 to promote ductal carcinoma in situ progression to invasive breast cancer by inducing epithelial-mesenchymal transition. Cancer cell. 2009 Sep 8;16(3):195–207.

[47] Paplomata E, O’Regan R. The PI3K/AKT/mTOR pathway in breast cancer: targets, trials and biomarkers. Therapeutic advances in medical oncology. 2014 Jul;6(4):154–66.

